# Sirtuin 1 modulation unlocks the therapeutic potential of sodium-glucose co-transporter 2 inhibitors (SGLT2i) in calcific aortic valve stenosis

**DOI:** 10.1101/2025.03.03.641336

**Authors:** Vincenza Valerio, Ilaria Massaiu, Matteo Franchi, Antoine Rimbert, Zoé Begué-Racapé, Mattia Chiesa, Veronika A. Myasoedova, Valentina Rusconi, Francesca Bertolini, Donato De Giorgi, Alice Bonomi, Sergio Pirola, Marco Zanobini, Paola Di Pietro, Albino Carrizzo, Michele Ciccarelli, Romain Capoulade, Stefano Genovese, Paolo Poggio

## Abstract

**Background:** Calcific aortic valve stenosis (AS) affects 3% of older adults and lacks medical treatment. The deacetylase Sirtuin 1 (SIRT1) could be involved in many pathways linked to AS progression. Sodium-glucose co-transporter 2 inhibitors (SGLT2i), glucose-lowering agents, have been shown to reduce cardiovascular events (likely *via* SIRT1), but their possible benefits in AS are unknown. Our study aims to uncover the role of SIRT1 in AS progression and assess the benefit of SGLT2i to slow down the aortic valve fibro-calcification processes.

**Methods:** RNA-seq data of human aortic valve specimens were collected from ARChS4 database. SIRT1 knockdown (SIRT1 KD) and overexpressing (SIRT1 Over) valve interstitial cells (VIC) were generated by CRISPR/Cas9. Real-time PCR, immunofluorescence, and calcification assays were used to characterized mutant VICs. Conditioned medium experiments were implemented to evaluate SGLT2i effect on cellular cross-talk and calcification. Diabetic patients’ data from the Lombardy regional healthcare database, treated with sulphonylureas (SU; no effect on SIRT1) and SGLT2i (acting on SIRT1), were selected and matched 1:1 by age, sex, and multisource comorbidity score. Cumulative incidence of hospitalization for non-rheumatic aortic valve disease was assessed by Kaplan-Meier and multivariable Cox proportional hazards models were used to estimate hazard ratios.

**Results:** RNA-seq showed that SIRT1 could be a master regulator of multiple AS-related pathways. Functional studies on mutant VICs revealed that SIRT1 directly regulates antioxidant processes, extracellular-matrix remodeling and calcification by modulating key transcription factors. Moreover, calcification assays further support this role, revealing an increased calcification in SIRT1 KD VICs and a concomitant decrease in VIC SIRT1 Over when compared to wild type. Then, exploring SGLT2i impact on calcification, we showed that VICs cultured in SGLT2i-treated-endothelial medium exhibited reduced calcification associated with endothelial-increased nitric oxide levels, while SIRT1 inhibition enhanced VIC calcification. The real-world data analysis revealed that SGLT2i-treated group had a lower incidence of hospitalized patients for non-rheumatic aortic valve disease compared to SU-treated group.

**Conclusions:** Our data identify SIRT1 as a key regulator of fibro-calcific processes in AS and suggest that SGLT2i may slow the aortic valve degeneration through SIRT1 modulation. These findings highlight SGLT2i as a promising therapeutic option for AS prevention and care.

**Clinical Perspectives:** *What is new?:*

- Sirtuin 1 (SIRT1) downregulation is linked to the progression of aortic stenosis (AS) pathological processes and plays a crucial role in mitigating oxidative stress, fibrosis, and calcification.
- Sodium-glucose co-transporter 2 inhibitor (SGLT2i) treatment of valve endothelial cells results in the secretion of protective factors that reduce valve interstitial cell calcification, highlighting the valuable role of endothelial health in preventing valve degeneration.
- Real-world data indicate that the use of SGLT2i is associated with a lower incidence of hospitalization rate for aortic valve disease as compared to sulfonylureas, suggesting a protective effect against AS in diabetic patients.

*What are the clinical implications?:*

- This study highlights the potential of restoring SIRT1 activity as a therapeutic strategy to mitigate pro-calcific and pro-fibrotic processes in AS.
- SGLT2i may offer a breakthrough therapeutic option for AS, a condition currently lacking effective pharmacological treatments, providing new hope for slowing disease progression and improving clinical outcomes.

## INTRODUCTION

Calcific aortic valve stenosis (AS) is a slowly progressing valve disease affecting approximately 2-7% of the population over 65^1^. This condition is marked by restricted leaflets mobility, a reduction in the aortic valve area, and a progressive thickening and calcification of the leaflets that ultimately leads to a significant obstruction of the left ventricular outflow tract, hindering the heart’s ability to function effectively^2^. With no pharmacological treatments currently available, surgical or transcatheter aortic valve replacement emerges as the only promising option^3^. A more profound grasp of the molecular underpinnings of AS is imperative to forge novel avenues for prevention and treatment opportunities, reducing the socio-economic burden currently imposed by its sequela.

AS is a complex process, and several cell types are pivotal in its progression. Mainly, valve interstitial cells (VICs) are the most studied cells in the pathology, given their crucial role in the aberrant production of extracellular matrix and calcium deposition. However, an equally key role is played by the thin valve endothelial cells (VECs) layer covering the cusps surface, orchestrating exchanges and communicating with the underlying VIC. Indeed, endothelial dysfunction is an inescapable first step in AS onset and development, as well as a significant driving force, which allows oxidized lipids to infiltrate the endothelium and thus attract inflammatory cells^4^. Based on this, the vital physiological and pathological interplay between these two cell types become clear, emphasizing the need for in-depth exploration, especially in the context of testing the effectiveness of new potential drug treatments.

Sirtuin 1 (SIRT1) is a deacetylase, known for its involvement in preventing senescence, cellular ageing, and endothelial dysfunction^5^. Its role is also crucial in regulating other cellular processes, including oxidative stress, inflammation, and vascular calcification, all of which involved in various cardiovascular diseases (CVD), including AS^6,7^. The observed down-regulation of SIRT1 in diseased aortic valve tissue^8^ suggests a potential link with AS, though this connection remains unexplored. Moreover, in the atherosclerosis contest, SIRT1 deficiency was associated with an increased expression of proprotein convertase subtilisin/kexin type 9 (PCSK9) gene, a fundamental regulator of cholesterol metabolism that has also been identified as a contributor of valve calcification^9,10^. In parallel, while glucose-lowering therapies, such as sodium-glucose co- transporter 2 inhibitors (SGLT2i), have shown promising cardiovascular benefits^11^, their potential impact on AS remains unmapped. Notably, SGLT2i has been shown to enhance SIRT1 cardioprotective effecte^12^, pointing to a possible therapeutic application against AS that merits further investigation. Given this context, it becomes intriguing to explore whether SIRT1 plays a role in the pathophysiological mechanisms of AS and whether SGLT2i could be proposed as a novel therapeutic strategy.

## MATERIAL AND METHODS

### Research oversight and data availability

RNA-sequencing data of aortic valve tissue samples were retrieved from the ARChS4 database^13^. A final dataset of 62 samples (26 healthy/non-calcified and 36 calcified valves) was analyzed. Additionally, a microarray-based gene expression dataset of 240 human AS specimens was retrieved from GEO (GSE102249).

Aortic valve specimen collection, for cell isolation from AS patients, was conducted in accordance with the principles outlined in the Declaration of Helsinki and approved by the Ethics Committee of the Centro Cardiologico Monzino (CCM 1068 – RE 4061). All participants provided written informed consent and RNA- sequencing data from wild type and SIRT1 mutant iVICs are available in the NCBI Sequence Read Archive (SRA) under BioProject ID PRJNA122727.

The analysis of genetic variants in the *SIRT1* locus, testing their associations with AS and related cardiovascular traits, were carried out using the largest genome-wide association meta-analysis available to date, which included 14,819 AS cases among 941,863 participants^14^. The screening for variants associated with expression levels of genes located in the *SIRT1* locus were performed using whole blood expression quantitative trait loci (eQTL) dataset (GTEX). The association of the top associated variant with multiple circulating biomarkers and cardiometabolic related factors was tested in the Pan-ancestry genetic analysis of the UK Biobank [released June 15, 2020].

The retrospective real-world study population analysis was based on Healthcare Utilization databases of Lombardy Region (Italy). Detailed descriptions of the regional databases and their use in studies involving patients with type 2 diabetes are reported elsewhere^15^.

### General material and methods

All material and methods for data analysis, cell processing, quantitative real-time reverse transcription PCR, bulk and single cell RNA-sequencing, enzyme-linked immunosorbent assays, capillary Western blotting, immunofluorescence staining and analysis, cell metabolic activity, reactive oxygen species measurement, calcification assay, secretome analysis, and nitric oxide evaluation are provided in the accompanying **Supplemental File**.

#### Statistical analysis

Data were analyzed using GraphPad Prism statistical software (version 7) or R or SAS V.9.4. Statistical comparisons between two groups were conducted using parametric Student t-test or unpaired non-parametric Mann-Whitney test, while comparisons between multiple groups were compared using one-way analysis of variance (ANOVA) with Tukey or Sidak or Dunnet post-hoc tests or two-way ANOVA with Tukey post-hoc test. Differences between groups were considered significant at p < 0.05 and all values are expressed as median with interquartile range (IQR).

Statistical comparisons between patients’ baseline characteristics of the two groups were conducted using standardized differences (SD) and SD <0.10 were considered negligible. Cumulative incidence of hospitalization for non-rheumatic aortic valve disease was estimated by using the Kaplan-Meier estimator, and the Log-rank test was used to assess differences between groups. Multivariable Cox proportional hazards models were used to estimate HR and 95% CI for risk of hospitalization for non-rheumatic aortic valve disease between patients who received SGLT2i compared with those who received SU. Covariates included various medications use such as statins, antihypertensives, antiplatelets, nitrates, antidepressants, nonsteroidal anti-inflammatory drugs, alpha-glucosidate inhibitors, thiazolidinediones., and respiratory drugs, as well as for the presence of renal diseases, respiratory conditions, and cancer. The assumption of proportionality of hazards were assessed using the Schoenfeld residuals. Stratified analyses were performed by sex and age (≤ 65 years vs > 65 years).

## RESULTS

### SIRT1 acts as a gatekeeper in AS pathophysiology

Many cellular and molecular processes are known to be involved in AS pathophysiology^16^. However, while the pathological mechanisms leading to valvular calcification were widely addressed, few studies have focused on the protective players^17^. In line with the European open data suggestions, publicly available RNA- seq data from non-calcified and calcified aortic valve tissues were collected and meta-analyzed to explore potential new candidates to fight AS degenerative disease.

We analyzed 62 human adult tricuspid aortic valves to shed light on key players associated with AS. Two clusters between AS (n = 36) and controls (CTRL, n = 26) specimens were highlighted from the multidimensional scaling (MDS) plot (**Figure 1A**) and 5,937 differentially expressed genes (adj p < 0.05) were identified between AS and CTRL, with 745 up-regulated (logFC > 1) and 395 down-regulated (logFC < -1) genes (**Figure 1B; Figure S2 and Table S7**). The top significantly regulated genes and the identified enriched functional pathways (**Figure 1C through 1H; Table S8**) are among the well-established hallmarks of AS^17^. In particular, our results highlight the positive regulation of several processes involved in cell activation (*e.g.*, migration, angiogenesis, interferon signaling, chemotaxis, and endothelial to mesenchymal transition), adaptive and innate immune response, extracellular matrix remodeling, signaling processes (*e.g.*, calcium, hippo, non-canonical Wnt, purinergic, and toll-like receptor signaling), cell death, and general processes (*e.g.*, platelet aggregation, tissue biomineralization, oxidative stress, and aging) in AS leaflets compared to controls. Moreover, other key biological mechanisms were found to be negatively regulated in AS specimens, such as fatty acid metabolism, mitochondrial protein, and ATP synthesis.

**Figure 1.**
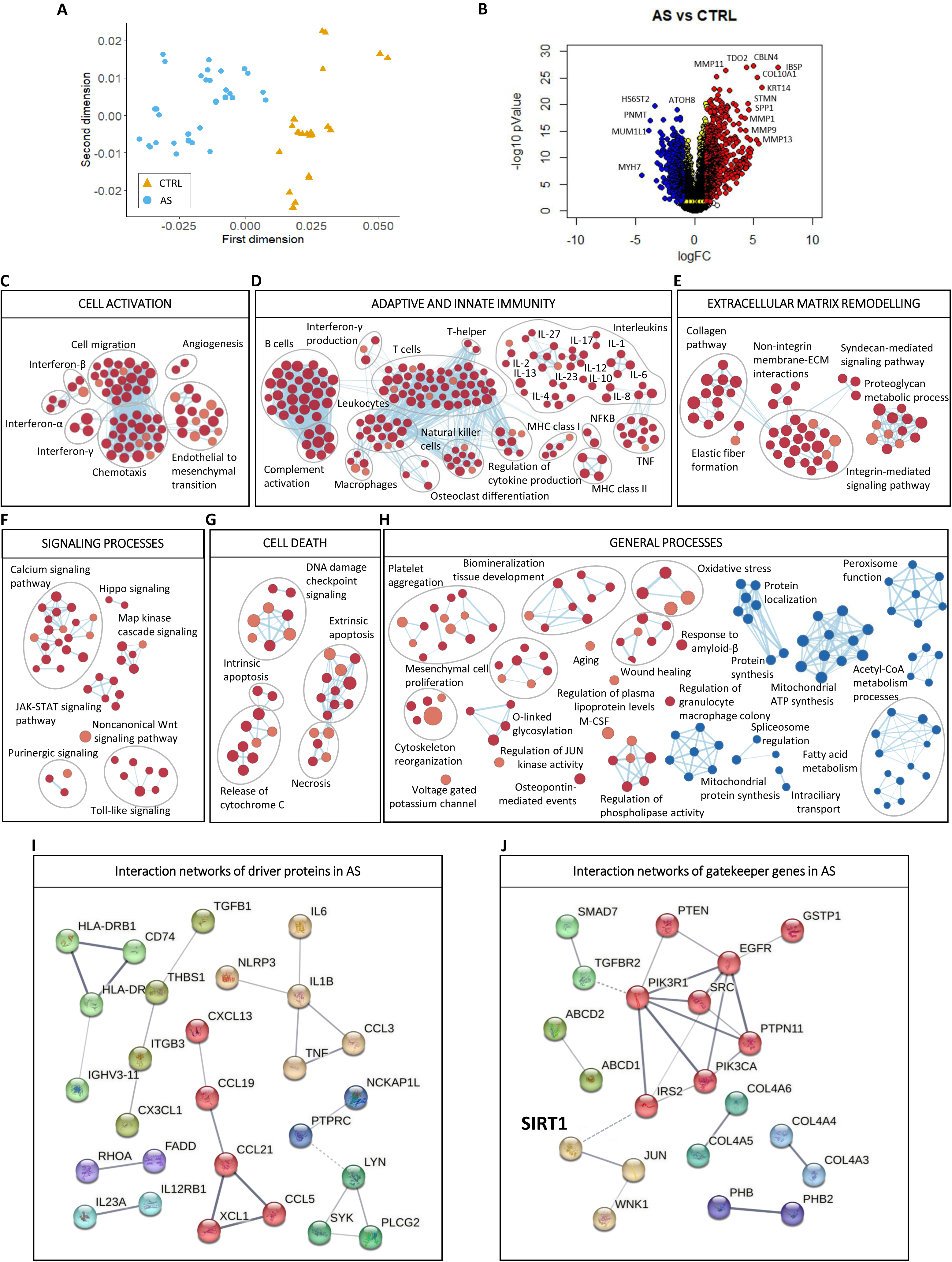
*SIRT1* as a gatekeeper in AS pathophysiology. **A**, Multidimensional scaling (MDS) plot of collected public RNA-seq data. Each dot represents one sample. Orange triangles: non-calcified subjects/controls (CTRL). Light blue circles: stenotic (AS) patients. **B**, Volcano plot showing the differential expression results of AS respect to CTRL. The genes with adjusted pValue > 0.05 are regarded as the differentially expressed genes (yellow dots) in AS compared to the CTRL group. In particular, red and blue dots denote significantly up-regulated (adjusted pValue < 0.05 and log(FC) ≥ 1) and down-regulated (adjusted pValue < 0.05 and log(FC) ≤ -1) genes in AS, respectively. The empty dots represent the unchanged genes. **C-H**, Functional pathways positively and negatively enriched in AS subjects. GSEA method was run and single pathways (nodes) represented through the Enrichment Map software were manually curated and clustered into six global functional networks (cell activation, adaptive and innate immunity, extracellular matrix remodelling, signaling processes, cell death, and general processes). The red and blue nodes represent the pathways positively and negatively enriched in AS samples (FDR qValue < 0.1), with a size proportional to the gene-set size. **I** and **J**, Interaction networks between the AS drivers and gatekeepers obtained from STRING database. The different nodes in the network represent the proteins while the network edges represent specific and meaningful protein-protein associations. The line thickness indicates the strength of data support. The solid lines represent the interactions between the proteins of the same cluster, while the dotted lines are for the interaction between the proteins of different clusters. Each identified cluster is shown through a different colour.

Selecting the differentially expressed genes with the highest degrees of connectivity (**Table S9**), we identified driver and gatekeeper proteins in AS pathophysiology (**Table S10**). Among the AS drivers, as recently highlighted^18^, pro-inflammatory cytokines can be noted, such as interleukin 6 (*IL6*), interleukin 1 beta (*IL1B*), tumor necrosis factor (*TNF*), and transforming growth factor beta 1 (*TGFB1*; **Figure 1I**). On the other hand, among the proteins with a potential protective role, the well-known deacetylase sirtuin 1 (*SIRT1*) stands out (**Figure 1J**).

As the role of SIRT1 in the development and progression of AS has been poorly explored to date, we turned our attention to clarify its impact on this complex pathology. Therefore, SIRT1 upstream modulators and downstream modulated players were investigated (**Figure S3A** and **S4A**). By experimental evidence^19^, protein kinase amp-activated catalytic subunit alpha 2 (*PRKAA2*), able to induce SIRT1 activation, was lowly expressed in AS compared to CTRL (p = 0.0032). Conversely, known suppressors of SIRT1 expression or activity^20,21^, such as *CD28*, interleukin 2 receptor subunit alpha and beta (*IL2RA/B*), cathepsin B (*CTSB*), retinoic acid inducible neural specific 1 (*BRINP1*), and cyclin-dependent kinase 5 (*CDK5*), were all up-regulated in AS (all p < 0.05; **Figure 2A** and **2B**). Considering transcription factors (TF), we found that the ones able to promote SIRT1 expression^22^ were down-regulated in AS, such as MYC proto-oncogene (*MYC*; p = 0.0011) and forkhead box O3 (*FOXO3*; p = 0.0052), while an inhibitor TF, hypermethylated in cancer 1 (*HIC1*), was up-regulated in AS (p = 7.7E-06, **Figure 2C**), resulting in a decreased expression of *SIRT1* in AS specimens (**Figure 2D**). These positive and negative associations were confirmed analyzing a public microarray-based expression dataset of 240 human stenotic valves. In addition, evaluating separately male and female patients (120 for each sex), we observed some sex-specific associations (**Figure S3B and S3C; Figure S4B and S4C**). Finally, analysis of the down-stream SIRT1 pathways, in accordance with few available studies^23,24^, we highlight the involvement of three critical pathways linked to AS pathophysiology (*e.g.*, antioxidant, fibrosis, and calcification processes; **Figure 2E through 2G**).

**Figure 2.**
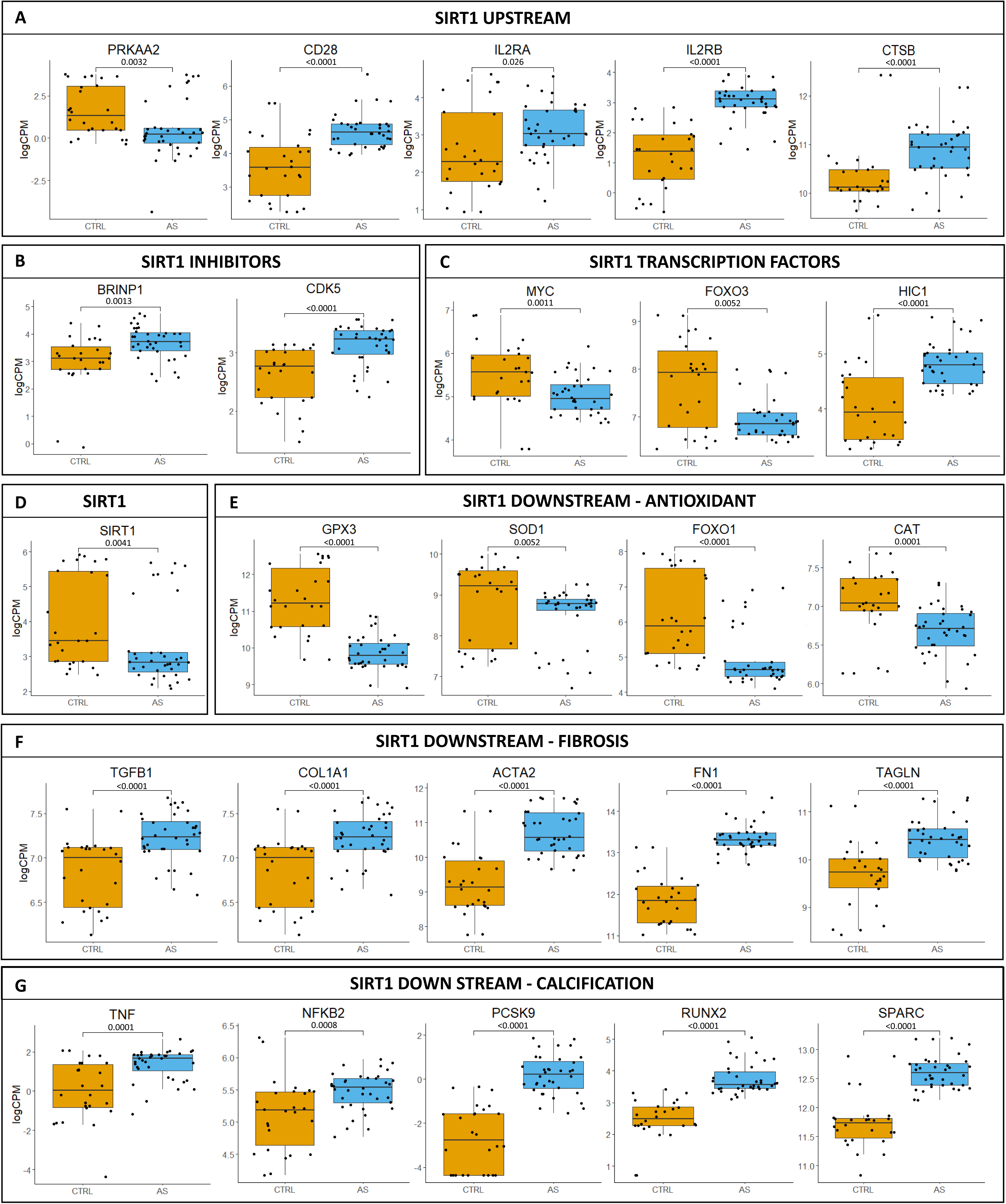
Dysregulation of *SIRT1* upstream modulators and downstream pathways in AS. Box and dot plots showing the logCPM values of relevant genes directly connected to SIRT1 for each sample grouped by condition (non-calcified subjects/controls (CTRL) in orange and stenotic (AS) patients in light blue). **A**, Expression levels in CTRL and AS samples of *PRKAA2*, *CD28*, *IL2RA*, *IL2RB*, and *CTSB* genes, connected to *SIRT1* at upstream level. **B**, Expression levels in CTRL and AS samples of *MYC*, *FOXO3*, and *HIC1* genes acting as transcription factors for *SIRT1*. **C**, Expression levels in CTRL and AS samples of *BRINPI* and *CDK5* genes, known as *SIRT1* inhibitors. **D**, *SIRT1* expression distribution in CTRL and AS samples. **E**, Expression distribution in CTRL and AS samples of *GPX3*, *SOD1*, *FOXO1*, and *CAT* genes, connected to *SIRT1* at downstream level and associated to antioxidant pathway. **F**, Expression distribution in CTRL and AS samples of *TGFBI*, *COL1A1*, *ACTA2*, *FN1*, and *TAGLN* genes, connected to *SIRT1* at downstream level and associated to fibrosis pathway. **G**, Expression distribution in CTRL and AS samples of *TNF*, *NFKB2*, *PCSK9*, *RUNX2*, and *SPARC* genes, connected to *SIRT1* at downstream level and associated to calcification pathway. Data are presented as median ± interquartile range with minimum and maximum values. The comparisons between CTRL and AS samples were performed using the non-parametric Mann- Whitney test. *** for pValues ≤ 0.001, ** for pValues ≤ 0.01, and * for pValues ≤ 0.05.

#### SIRT1 overexpression modulates transcription factors involved in anti-oxidant, pro-fibrotic, and pro-calcific processes

Building on our previous findings and on previous evidence from literature^8^, we investigated the causative role of SIRT1 in AS pathophysiological processes. To achieve this, immortalized VICs (iVIC) were genetically modified to generate stable *in vitro* models with down-regulated or up-regulated *SIRT1* gene expression (over- and KD-iVICs, respectively) and functionally characterized (**Figure 3A through 3D; Figure S5**).

**Figure 3.**
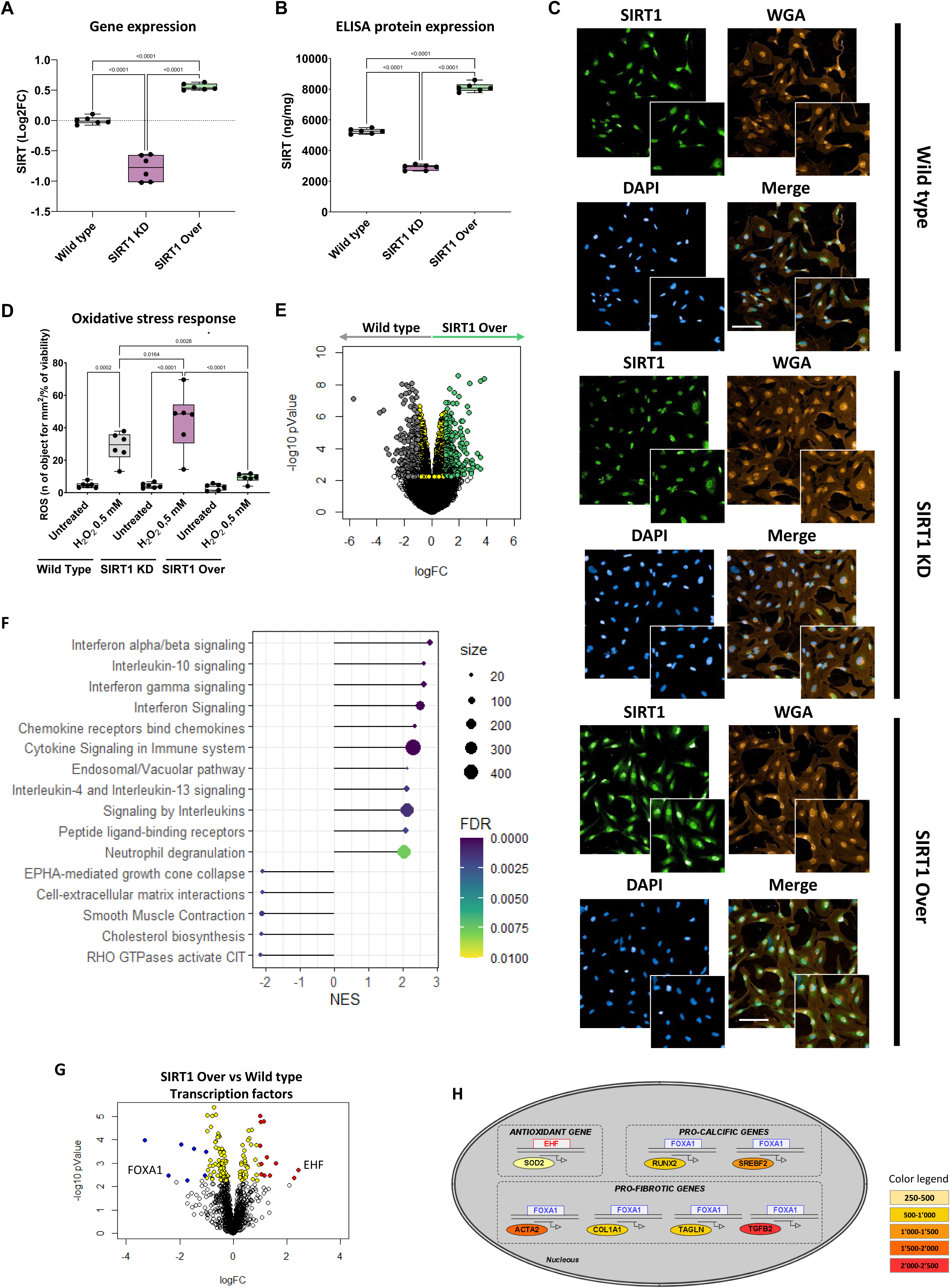
*SIRT1* overexpression modulates key transcription factors and pathways in AS pathogenesis. Box and whisker plots representing gene expression (**A**) and protein expression (**B**) levels of SIRT1 in Wild Type (grey; n = 6), SIRT1 KD (purple; n = 6) and SIRT1 Over (light green; n = 6) cells. Represented pValue are multiple comparison after ANOVA with Tukey correction. **C**, Immunofluorescence staining showing SIRT1 (*green)* expression and localization in Wild Type, SIRT1 KD and SIRT1 Over stenotic iVICs. Membrane were visualized with WGA (*orange*) and nuclei with DAPI (*blue*). Scale bar: 100 µM. **D**, Box and Whisker plot representing oxidative stress response in term of ROS production in Wild type (n = 6), SIRT1 KD (n = 6) and SIRT1 Over (n = 6) cells, after treatment with H_2_0_2_. Represented pValues are multiple comparison after ANOVA with Sidak correction. Data are presented as median ± interquartile range with minimum and maximum values. **E**, Volcano plot showing the results of the differential expression analysis of SIRT1 Over compared to Wild Type subjects. The genes with adjusted pValue < 0.05 are considered differentially expressed (DEGs, yellow genes). Grey and green dots denote significantly up- regulated genes in Wild Type (log(FC) ≤ 1) and SIRT1 Over (log(FC) ≥ 1) samples, respectively. The empty dots represent the unchanged genes. **F**, Lollipop plot of the positively (normalized enrichment score (NES) > 0) and negatively (NES < 0) enrichment pathways in SIRT1 Over samples respect to WT. The gradient colour represents the FDR value and the dot size is proportional to the leading-edge number. **G**, Volcano plot representing exclusively the differentially expressed transcription factors in SIRT1 Over respect to WT samples (│log(FC)│ ≥ 1 and adjusted pValue > 0.05). In particular, red and blue dots denote significantly up- regulated (adjusted pValue < 0.05 and log(FC) ≥ 1) and down-regulated (adjusted pValue < 0.05 and log(FC) ≤ -1) transcription factor in SIRT1 Over, respectively. The empty dots represent the unchanged transcription factors. **H**, Representation of the relation of *EHF* and *FOXA1* transcription factors (square) with key target genes (circles) driving the antioxidant (*SOD2*), pro-calcific (*RUNX2* and *SREBF2*), and pro-fibrotic (*ACTA2*, *COL1A1*, *TAGLN*, and *TGFB2*) processes. *ZFY* (depicted in red) is an up-regulated transcription factor in SIRT1 Over samples, while *FOXA1* (depicted in blue) is down-regulated. A color gradation representing the binding strength of the transcription factor to the target gene is used and indicated in the legend.

RNA sequencing was performed to elucidate the differences between mutant iVICs at steady-state (**Figure S6)**. This analysis revealed 225 up-regulated and 228 down-regulated genes in SIRT1 Over iVICs compared to wild type iVICs (**Figure 3E; Figure S7A and Table S11**), whereas only 4 down-regulated genes were identified in SIRT1 KD iVICs compared to wild type (**Figure S7B and S8, and Table S11**). In addition, as shown from the differential analysis, multiple regulated pathways emerged in SIRT1 Over iVICs (**Figure 3F and Table S12**). We observed an up-regulation of pathways related to interferon alpha/beta signaling, which intriguingly coexisted with anti-inflammatory interleukin 10, 4, and 13 signaling in SIRT1 Over iVICs. Furthermore, pathways associated with extracellular matrix remodeling and myofibroblastic activation were found to be down-regulated by SIRT1 overexpression.

Based on an integrated analysis of the collected public RNA-seq data of CTRL and AS aortic valve tissues and our RNA-seq data from WT and SIRT1 Over iVICs, we extracted 438 genes linked to AS pathology and consistently regulated by SIRT1 (**Table S13**). Among these genes, we found *ADAMTS7*, *LMO4*, and *TMEM44* which were recently identified as genes associated with AS in the latest genome-wide association study (GWAS) meta-analysis^14^ (**Figure S9**).

Focusing on the SIRT1-regulated TFs (**Figure 3G; Table S14**), the top up-and down-regulated ones, with known and validated target genes, were ETS homologous factor (*EHF*) and forkhead box A1 (*FOXA1*), respectively. Specifically, EHF is responsible for the antioxidant response, regulating the *SOD2* gene, which is significantly up-regulated in SIRT1 Over cells. In contrast, *FOXA1* is up-stream of pro-fibrotic (*ACTA2*, *COL1A1*, *TAGLN*, *TGFB2*) and pro-calcific genes (*RUNX2*), all significantly down-regulated in SIRT1 Over cells beside *RUNX2* (**Figure 3H; Figure S10**). Remarkably, *FOXA1* can also regulate the expression of SREBF2, which, in turn induces PCSK9 expression, which is acknowledged for its role in aortic valve calcification^25^. Therefore, our data shows that regulating SIRT1 may be functional to protect VICs from processes crucial in AS pathogenesis, such as oxidative stress, fibrosis, and calcification.

#### SIRT1 as a shield against fibrosis and calcification in AS

Based on evidences that SIRT1 can attenuate cardiac fibrosis^26^ and supported by the RNA-sequencing data from our mutant cells, we first assessed the expression of various fibrosis-related genes in SIRT1 KD and Over iVICs. We noted that the expression of *ACTA2*, *COL1A1*, *FN1*, *BGN*, *TAGLN*, and *TGFβ2* was significantly reduced in SIRT1 Over iVICs when compared to wild type and SIRT1 KD iVICs (**Figure 4A through 4F; Figure S11A**). At the protein level, we showed a significant reduction of αSMA in SIRT1 Over cells both in the absence and in the presence of a pro-fibrotic stimulus (**Figure 4G, 4I, and 4K; Figure S11B and S11D**). Likewise, collagen levels decreased in SIRT1 Over cells and increased in SIRT1 KD cells when compared to WT (**Figure 4H, 4J, and 4L; Figure S11C and S11E**). By examining the calcification process in the mutant cells, we observed an upward trend in RUNX2 expression, as well as a significant increase in TNF and PCSK9 expression in SIRT1 KD iVICs (**Figure 5A through 5C)**. Given the involvement of PCSK9 in AS calcification processes^10^, we assessed its protein levels in mutant iVICs. Our findings revealed a significant intracellular and extracellular increment of PCSK9 in SIRT1 KD iVICs, alongside with a significant decrease in SIRT1 Over cells (**Figure 5C and 5D**). Based on these data, we investigated the calcification potential of our mutant cells. In this regard, we found that iVICs SIRT1 KD exhibited a significant increase in calcification with a concomitant significant reduction in SIRT1 Over iVICs (**Figure 5F and 5G**). To gain deeper insights into the role of SIRT1 in calcification and its connection with PCSK9, we generated iVICs overexpressing PCSK9 (**Figure S12**). After gene and protein expression validation, we confirmed that these cells exhibited a higher propensity for calcification compared to the wild type (**Figure 5H and 5I**).

**Figure 4.**
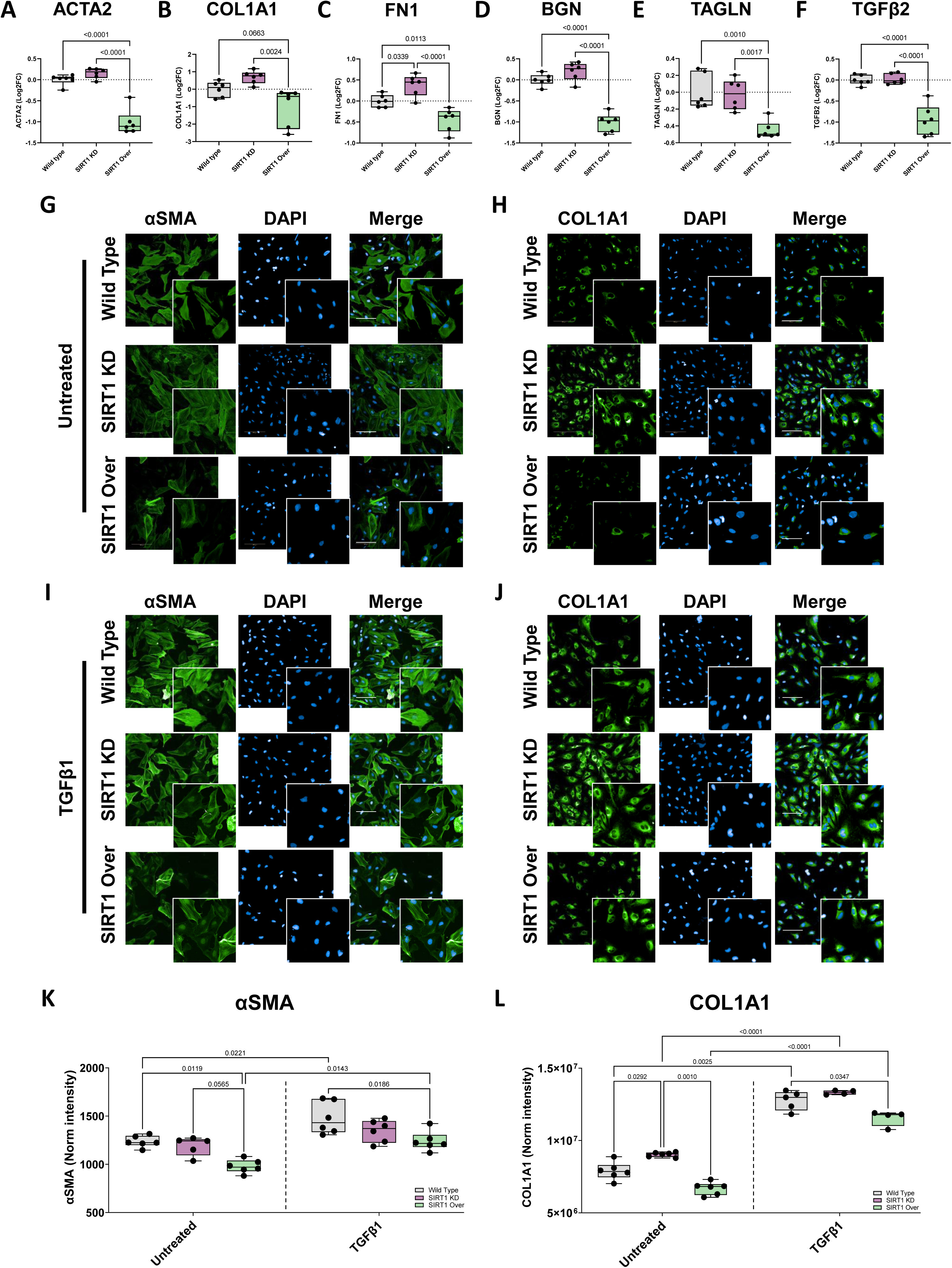
Protective role of SIRT1 towards fibrosis. Box and whisker plots representing gene expression of (**A**) *ACTA2*, (**B**) *COL1A1*, (**C**) *FN1*, (**D**) *BGN*, (**E**) *TAGLN*, (**F**) *TGFβ2* in Wild Type (grey; n = 6), SIRT1 KD (purple; n = 6) and SIRT1 Over (light green; n = 6) cells. Immunofluorescence staining showing αSMA (green) expression and localization in Wild Type, SIRT1 KD and SIRT1 Over stenotic iVICs in absence of treatment (Untreated) (**G**) and after treatment with TGFβ1 (**I**). Nuclei were visualized with DAPI (blue). Scale bar: 100 µM. Immunofluorescence staining showing COL1A1 (green) expression and localization in Wild Type, SIRT1 KD and SIRT1 Over stenotic iVICs in absence of treatment (Untreated) (**H**) and after treatment with TGFβ1 (**J**). Nuclei were visualized with DAPI (blue). Scale bar: 100 µM. Box and Whisker plot showing quantification of αSMA (**K**) and COL1A1 in iVICs Wild Type (n = 6) SIRT1 KD (purple; n = 6) and SIRT1 Over (light green; n = 6) in absence of treatment (Untreated) and after treatment with TGFβ1. Data are presented as median ± interquartile range with minimum and maximum values. All represented pValues are multiple comparison after ANOVA with Tukey correction.

**Figure 5.**
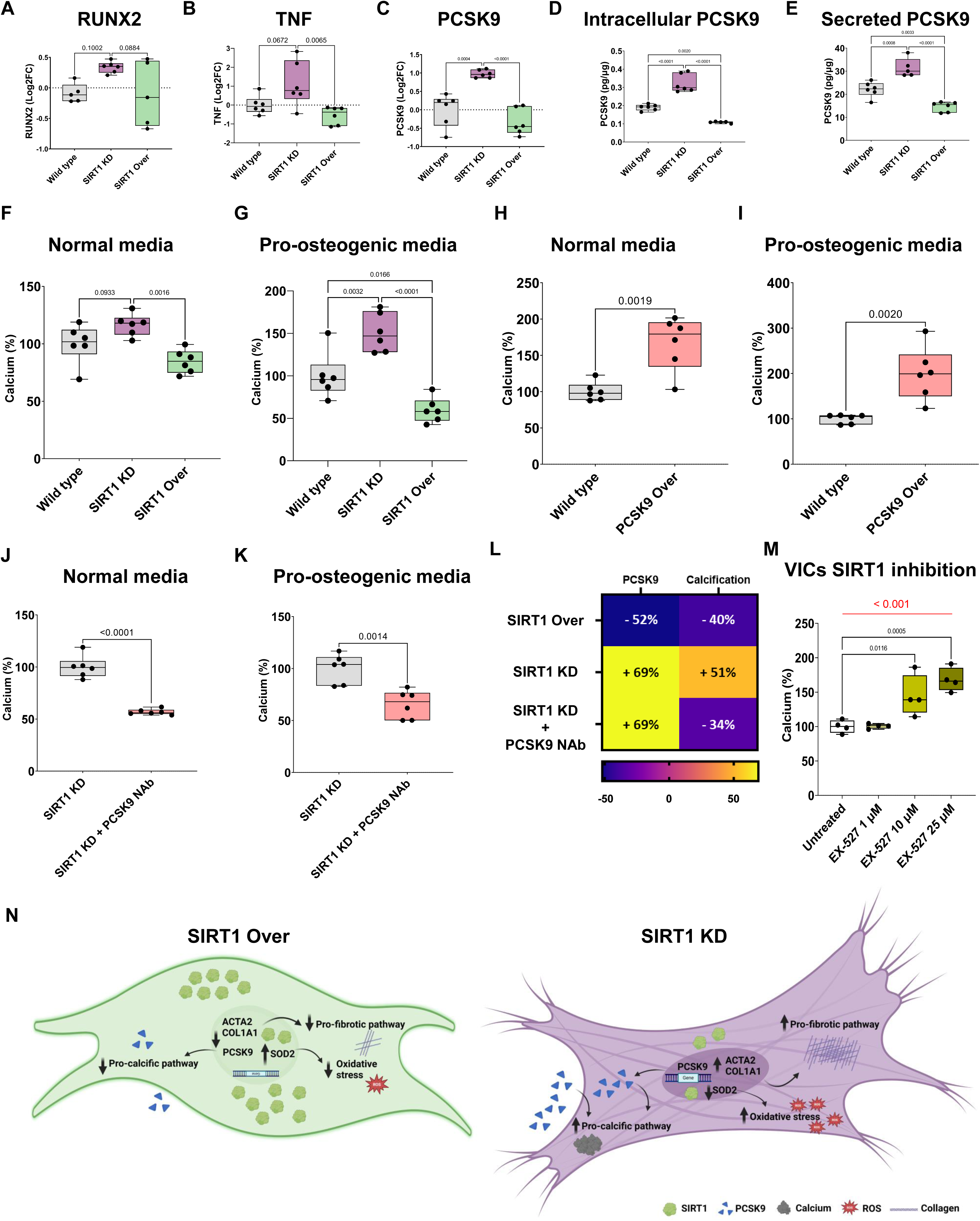
Protective role of SIRT1 towards calcification. Box and whisker plots representing gene expression of (**A**) *RUNX2*, (**B**) *TNF* and (**C**) *PCSK9* in Wild Type (grey; n = 6), SIRT1 KD (purple; n = 6) and SIRT1 Over (light green; n = 6) cells. Represented pValues are multiple comparison after ANOVA with Tukey correction. Box and whisker plots representing intracellular expression (**D**) and secreted levels (**E**) of *PCSK9* in Wild Type (grey; n = 6), SIRT1 KD (purple; n = 6) and SIRT1 Over (light green; n = 6) cells. Represented pValues are multiple comparison after ANOVA with Tukey correction. Box and whisker plots representing calcium percentage in Normal (**F**) and Pro-osteogenic media (**G**) in in Wild Type (grey; n = 6), SIRT1 KD (purple; n = 6) and SIRT1 Over (light green; n = 6) cells. Represented pValues are multiple comparison after ANOVA with Tukey correction. Box and whisker plots representing calcium percentage in Normal (**H**) and Pro-osteogenic media (**I**) in Wild Type (grey; n = 6) and PCSK9 Over (pink; n = 6) cells. Box and whisker plots representing calcium percentage in Normal (**J**) and Pro-osteogenic media (**K**) in SIRT1 KD without (grey n = 6) and with neutralizing antibody against PCSK9 (pink n = 6) treatment. Represented pValue is Student t-test. (**L**) Heatmap representing calcification percentage related to PCSK9 levels in SIRT1 Over, SIRT1 KD and SIRT1 KD treated with Neutralizing antibody against PCSK9. **M**, Box and whisker plot representing calcium percentage in stenotic primary VICs treated with different concentration of EX-527. p < 0.001 represent T for Trend (Red line); represented pValues are multiple comparison after ANOVA with Dunnet correction (Black lines). All data are presented as median ± interquartile range with minimum and maximum values. **N**, representative cartoon that summarizes the cellular changes observed in previous data Particularly, the green cell (SIRT1 OVER) shows that SIRT1 overexpression inhibits pro-fibrotic and pro-calcific pathways, reducing oxidative stress and ultimately preventing calcification. In contrast, the purple cell (SIRT1 KD) illustrates that SIRT1 knockdown activates these pathways, increasing oxidative stress and leading to enhanced calcification.

Furthermore, by blocking PCSK9 with a specific neutralizing antibody in SIRT1 KD iVICs, which exhibited elevated PCSK9 levels and higher calcification potential, we revealed that this inhibition led to a substantial reduction in calcification, thus, reinforcing the link between SIRT1 modulation and PCSK9 regulation (**Figure 5J through L**). Finally, we validated the inverse causal link between SIRT1 activity and calcification in primary stenotic VICs using a known SIRT1 inhibitor, namely EX-527 (**Figure 5M and 5N**).

### SIRT1 as a player in blood pressure and SGLT2i effects

Based on our previous results, we investigated the genetic association between *SIRT1* and AS. Utilizing the largest GWAS published to date, we did not identify any variants in the locus significantly associated with AS^14^ (**Figure S13**). However, upon querying the cardiovascular disease knowledge portal, we discovered that common variants in the *SIRT1* locus have been associated with several cardiovascular related traits, with the most significant signals for systolic blood pressure (SBP) and pulse pressure (**Figure 6A**).

**Figure 6.**
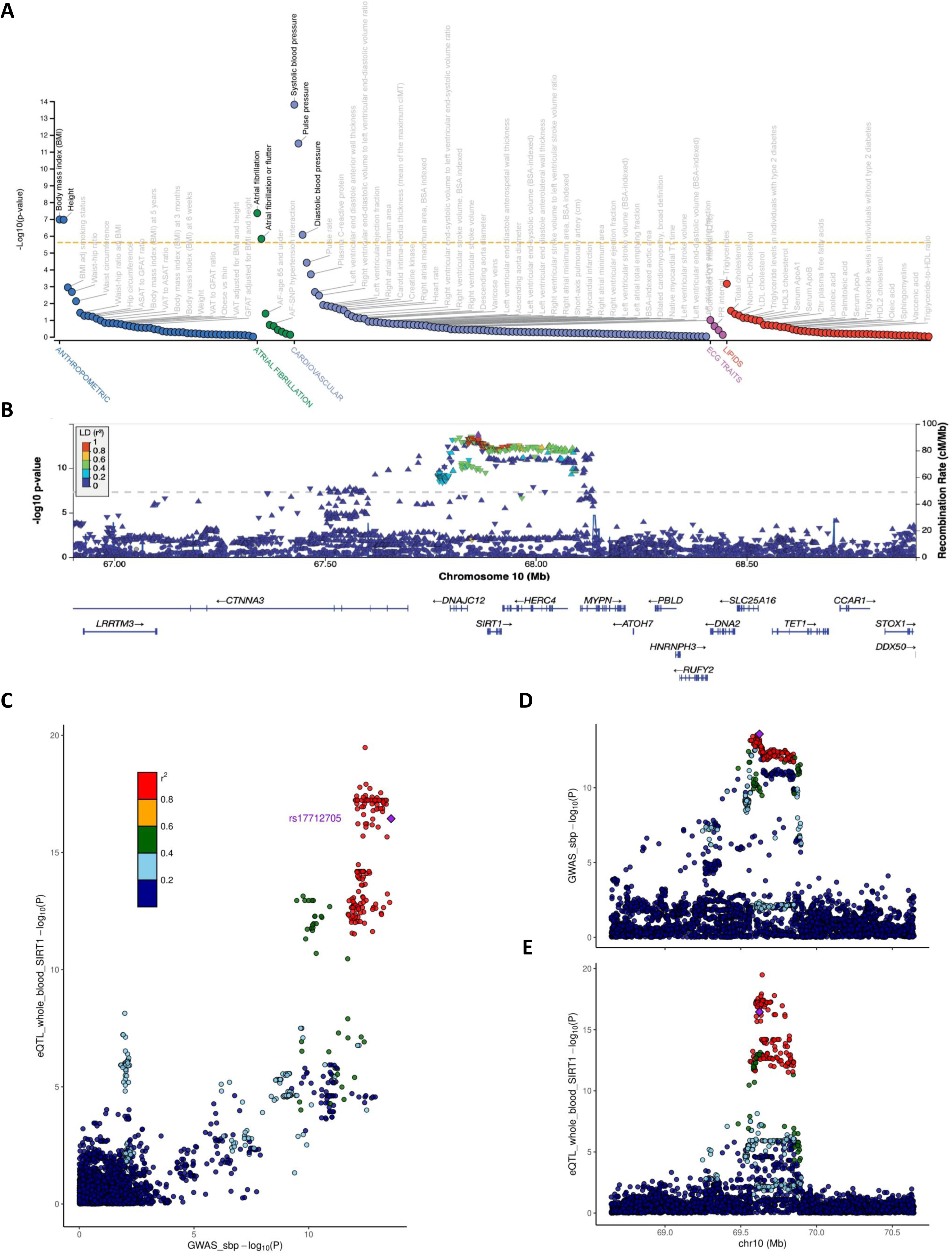
Genetic variants in SIRT1 region and their association with systolic blood pressure. **A**, Figure illustrating the associations between common genetic variants in the SIRT1 gene region and cardiovascular traits, as derived from data available in the Cardiovascular Disease Knowledge Portal (https://cvd.hugeamp.org/gene.html?gene=SIRT1). **B**, Locus zoom of genetic associations of variants in the SIRT1 gene locus (10q21.3) with systolic blood pressure. Data were extracted from the genome-wide association meta-analysis published by Keaton et al. 2024^27^ (n=1,028,980 individuals). Each variant identified in the 1000 genomes project is represented by a triangle, with color representing linkage disequilibrium (LD) with the lead SNP (rs17712705, purple diamond). **C**, Colocalization between SIRT1 locus in systolic blood pressure GWAS by Nikpay et al.^50^ and SIRT1 eQTL in whole blood. The eQTL pvalues were extracted from GTEx (n = 335 individuals). The GWAS pvalues were extracted from Nikpay et al.^50^ (n case = 60,801 and n control = 123,504 individuals). Respective Manhattan plots are presented in panel (**D**) and (**E**). Each variant identified in the 1000 genomes project is represented by a dot, with colour representing linkage disequilibrium (LD) with the lead SNP (purple diamond).

To further verify this finding, we screened the *SIRT1* locus for common genetic variants associated with SBP in the largest meta-analysis published to date^27^. Indeed, we found significant genetic associations of the top associated SNP rs17712705 with SBP (p = 2.55e-14) (**Figure 6B**). Notably, the associated variants in the *locus* encompass four genes: *DNAJC12*, *SIRT1*, *HERC4*, and *MYPN* (**Figure 6B**).

To provide additional information on the gene associated with SBP within the *locus*, we screened for variants associated with both SBP and the expression levels of nearby genes, assessed by expression quantitative trait loci (eQTL) dataset from GTEx (whole blood). Our analysis revealed that genetic variants associated with SBP are also significantly associated with changes in *SIRT1* gene expression, whereas they do not affect gene expression of *DNAJC12*, *HERC4*, and *MYPN* (**Figure 6C**; **Figure S14**). This data suggests that variants in the locus affect *SIRT1* gene expression, which is likely involved in SBP changes.

To gain further insights into phenotypic associations with *SIRT1*, we tested the association of rs17712705 with multiple circulating biomarkers and cardiometabolic-related traits (n = 150, **Table S4**) in the UK Biobank (∼500,000 participants) (**Figure S15**). We observed that rs17712705, associated with SBP and *SIRT1* gene expression, is significantly associated with traits related to kidney function (urate, urea, cystatin c, creatinine), blood pressure (SBP and pulse pressure) and glucose plasma levels.

Interestingly, we noted similarities in the pleiotropic effect of sodium-glucose cotransporter 2 inhibitors (SGLT2i) that influence blood glucose by reducing renal glucose reabsorption, affect blood pressure through natriuresis, and impact renal function by reducing intraglomerular pressure^28,29^. We therefore hypothesized that SIRT1 influence on SBP, which is one of them most critical AS risk factors, might be involved in or linked to SGLT2i functions.

### SGLT2i promotes valve endothelial cell-mediated reduction of valve interstitial cells calcification

Given the cardioprotective effects of SGLT2i and their role in enhancing SIRT1 activation^12^, we sought to evaluate their efficacy in aortic valve calcification taking into account the complexity of the valve structure and the natural cross-talk between the valve endothelial monolayer and the underlying VICs (**Figure 7A**).

**Figure 7.**
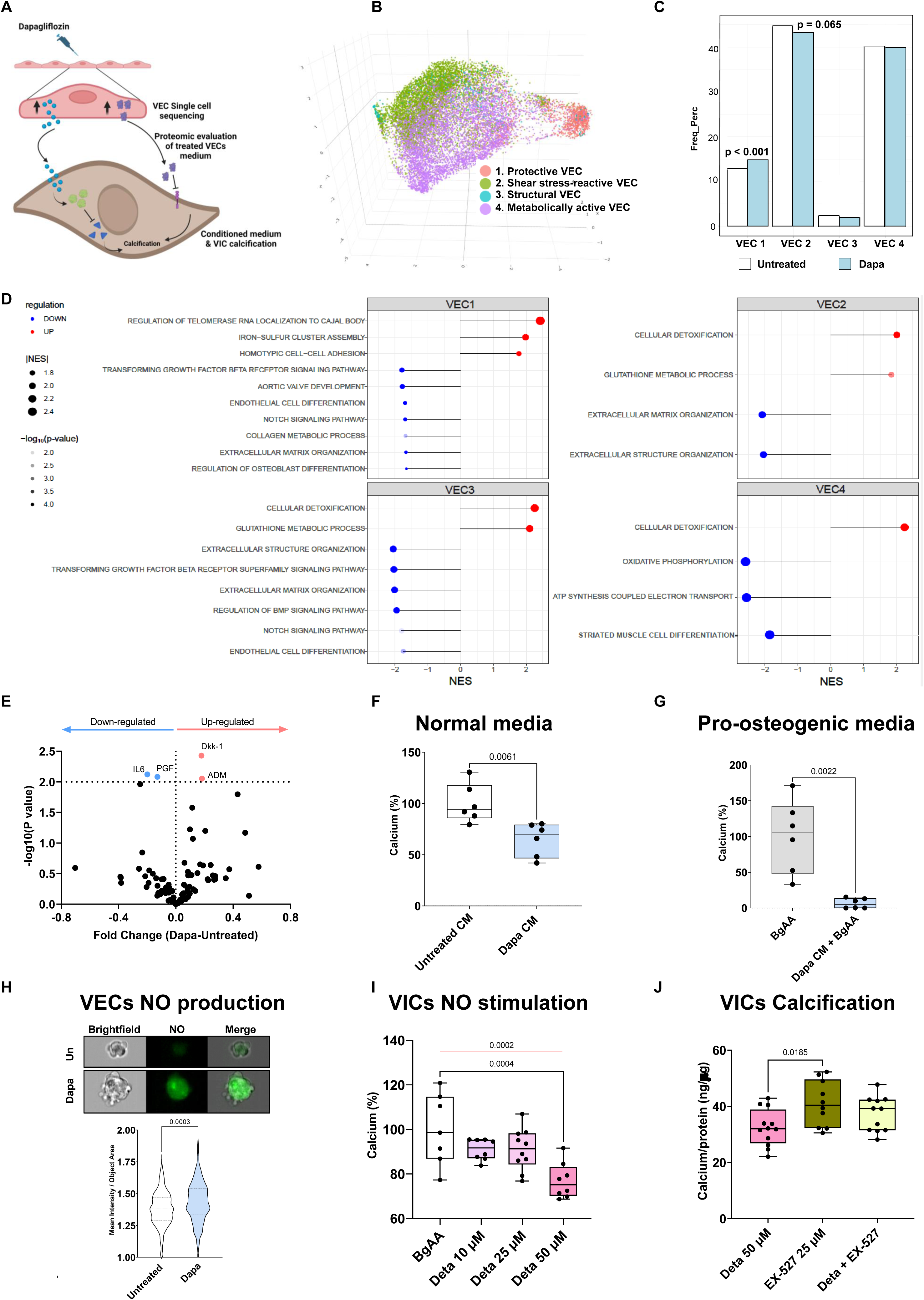
Beneficial effects of SGLT2i on protective cellular cross-talk against calcification. **A**, Schematic representation of the experimental workflow investigating the effects of Dapagliflozin on VECs and consequently VICs calcification. Dapagliflozin treatment is administered to VECs, which are then analyzed using single-cell sequencing to identify key populations. The conditioned medium from dapagliflozin-treated VECs is subjected to proteomic evaluation to identify secreted factors. These conditioned medium are then applied to VICs leading to the observation of reduced calcification. Arrows indicate the progression of treatment and analysis, with calcification inhibition as the final observed outcome. **B**, Tri-dimensional scatterplot of UMAP highlighting the inferred cell-types. Each dot represents a cell while colours refer to VEC subtypes: protective (red, VEC1), shear-stress reactive (green, VEC2), structural (cyan, VEC3), and metabolically active (purple, VEC4) VEC. **C**, Histogram showing the frequency of cells belonging to each cluster, considering Untreated (whit bars) and treated with Dapagliflozin (light blue) samples separately. Represented pValue is Chi-squared proportion test. **D**, Lollipop plot showing the significant pathways positively (red dots) or negatively (blue dots) associated to Dapagliflozin treatment in each VEC subtype. The size of dots is related to the absolute value of the Normalized Enrichment Score (NES) while the transparency refers to the significance; the darker the color the more significant the association. **E**, Volcano plot summarizing the statistical analysis of secreted proteins, assessed by Olink technology. Each dot is a protein while x- and y - axis represent the fold changes and the pvalue in logarithmic scale, respectively. Red and blue dots are protein up- and down- regulated when compare patient with and without Dapagliflozin treatment, respectively. **F**, Box and whisker plot representing calcium percentage in stenotic primary VICs treated with VECs conditioned medium in absence of treatment (white; n = 6) and after treatment with dapagliflozin (light blue; n = 6). Represented pValue is Student t-test. **G**, Box and whisker plot representing calcium percentage in stenotic primary VICs treated with VECs conditioned medium treated with BgAA (grey) and BgAA supplemented with Dapagliflozin (light blue). Represented pValue is non- parametric Mann-Whitney tests. **H**, Image flow cytometry analysis and violin plot of NO levels in primary VECs Untreated (Un; white) and treated with Dapagliflozin (Dapa; light blue). Magnification of images: 40x. Represented pValue is Student t-test. **I**, Box and whisker plot representing calcium percentage in stenotic primary VICs treated with BgAA (n = 6) and bgAA supplemented with different concentrations of Detanonate (deta; n = 6). p < 0.0002 represent T for Trend (Red line); other represented pValues are multiple comparison after ANOVA with Dunnet correction (Black lines). **J**, Box and whisker plot representing calcium production in stenotic primary VICs treated with detanonate (deta; n = 6; pink), EX-527 (green; n = 6) and Detanonate supplemented with EX-527 (Deta + Ex-527 n = 6). Represented pValues are multiple comparison after ANOVA with Tuckey correction. All data are presented as median ± interquartile range with minimum and maximum values.

Valve endothelial cells (VEC) were subjected to single-cell sequencing and 4 different clusters were identified, based on a previous study^30^: protective (VEC 1), shear stress-reactive (VEC 2), structural (VEC 3), and metabolically active (VEC 4) VECs (**Figure 7B**). Notably, after treatment with dapagliflozin, we observed a significant increase in the cell count of VEC 1 and a concomitant decreasing trend in VEC 2 number (**Figure 7C**). Moreover, the transcriptomes of each cell cluster were differently dis-regulated by the drug treatment (**Figure S16**), allowing the identification of enriched pathways specific to each cell type. Notably, in VEC 1 we observed up-regulation of pathways related to telomere maintenance and enhanced resistance to oxidative stress. At the same time, several processes associated with the pathophysiology of aortic valve stenosis, were down-regulated, including TGF-b and Notch signaling, endothelial differentiation and processes linked to fibrosis and calcification (**Figure 7D**, upper left panel). Up-regulated pathways related to cell detoxification and glutathione metabolism were identified in VECs 2, 3, and 4. Simultaneously, pathways related to extracellular matrix organization were found to be down-regulated in VEC 2 and VEC 3, with the addition of TGF-b and Notch signaling pathways specifically down-regulated in VEC 3. In contrast, VEC 4 showed down-regulation of ATP synthesis pathways and oxidative phosphorylation pathways (**Figure 7D**).

In the context of cellular cross-talk, we performed a proteomic analysis on the conditioned medium of dapagliflozin-treated VECs. This analysis revealed a down-regulation of pathology-aggravating factors, including Interleukin-6 (IL6)^31^ and placenta growth factors (PGF)^32^, in the medium of treated cells compared to untreated controls. Conversely, we observed an up-regulation of factors potentially protective against calcification, such as Dickkopf 1^33^ (DKK1) and Adrenomedullin^34^ (ADM) (**Figure 7E**).

Based on these finding, the conditioned medium of VECs was administered to VIC, showing a significant decrease in calcification (**Figure 7F and 7G**). Given that SGLT2i up-regulates ADM, also known to induce nitric oxide (NO) production^35^, and to mitigate endothelial dysfunction *via* the SIRT1-eNOS axis, we assessed NO production. We observed a significant increase in NO levels in dapagliflozin-treated VECs (**Figure 7H**). To further investigate the role of the SIRT1-NO axis in reducing VIC calcification, we treated VIC with Deta Nonoate, a NO donor, and EX-527, the SIRT1 inhibitor. Our results showed that NO effectively reduces VICs calcification, whereas EX-527 abrogates this beneficial effect, thereby confirming the connection between SGLT2i, NO pathway and SIRT1 (**Figure 7I and 7J**; **Figure S17**).

### Real-world evidence of SGLT2i protective role against aortic valve degeneration

Based on the *in vitro* results showing the encouraging efficacy of SGLT2i in reducing calcification of VICs through cross-talk with VECs, we sought to investigate further the potential protective role of SGLT2i on valve degeneration. To this end, we conducted a large, retrospective, real-world population-based analysis. Among the 233,695 eligible patients who started second-line therapy with either sulphonylureas (SU; n = 192,007) or SGLT2i (n = 41,668) between 2008 and 2020, a total of 30,172 patients treated with SGLT2i were 1:1 matched with as many patients treated with SU by date of start of second line therapy, age, sex, and multisource comorbidity score (MCS; **Figure 8A; Table S15**).

**Figure 8.**
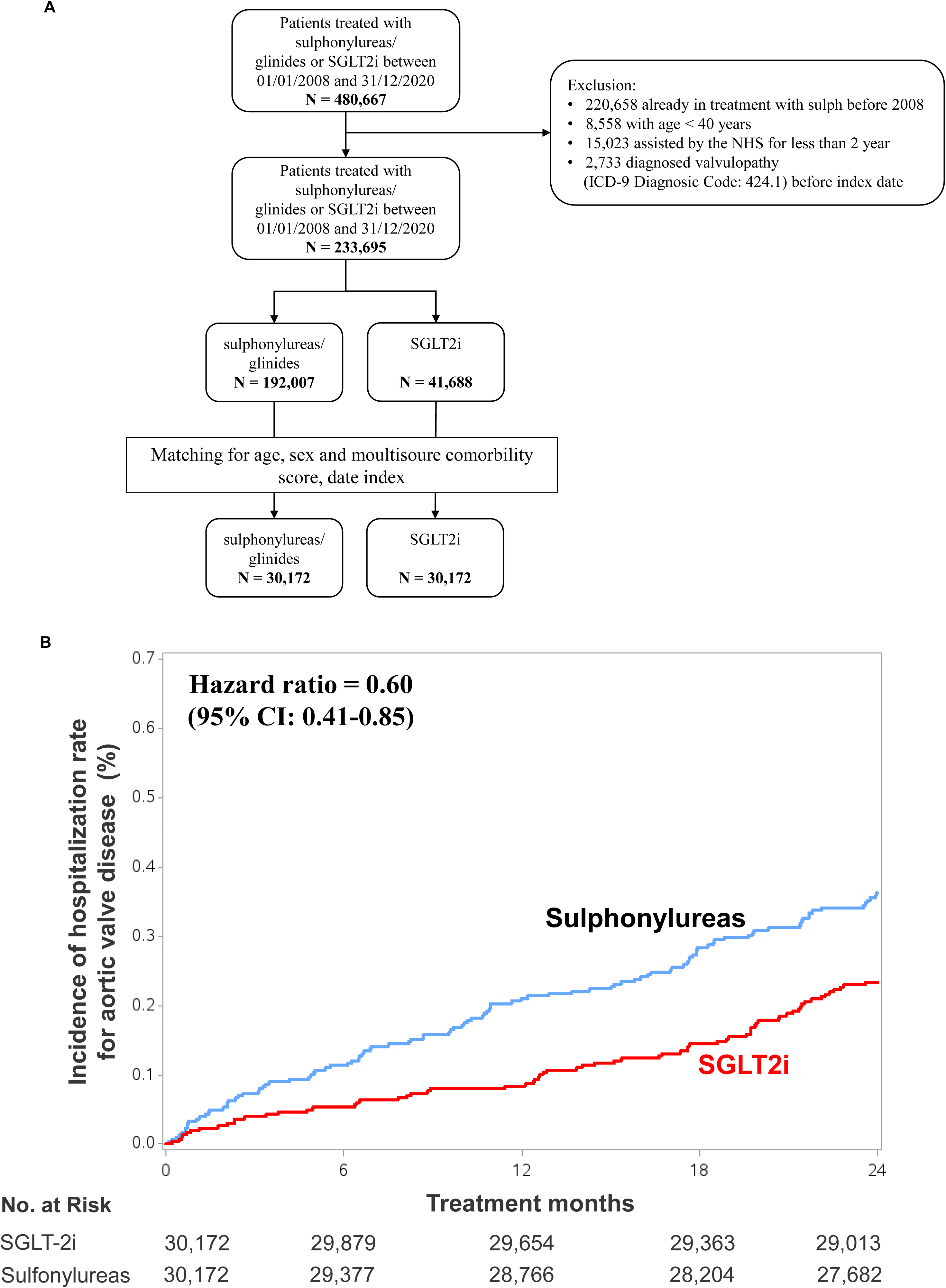
Real-world evidence of SGLT2i protection from hospitalization for non-rheumatic aortic valve disease. **A**, Flow-chart represented the inclusion and exclusion criteria, as well as the patient matching process employed in the study. **B**, Kaplan-Meier curve representing the incidence of hospitalization for non-rheumatic aortic valve disease in SGLT2i-treated patient (red line) vs SU-treated patients (blue line) over a 24-minth follow-up period.

Kaplan-Meier curves were performed to compare the cumulative incidence of hospitalization for aortic valve disease between patients treated with SU and SGLT2i over a 24-month follow-up period. During the follow-up, the percentage of the hospitalizations remained consistently higher among SU group compared to SGLT2i group, reaching 105 and 69 events, respectively (0.35% *vs.* 0.23%; p = 0.0063). The adjusted hazard ratio was 0.60 (95% CI 0.41–0.85), indicating a 40% lower hospitalization rate for aortic valve disease diagnosis in the SGLT2i group relative to those that received SU (**Figure 8B**). Additional analyses were performed by dividing the patient cohort into those under and over 65 years of age, as well as by sex-separating male and female patients (**Figure S18**). The SGLT2i-treated group consistently demonstrated a lower percentage of patients hospitalized for non-rheumatic aortic valve disease over time compared to SU-treated group, highlighting the potential cardioprotective effects of SGLT2 inhibitors in this population. All the calculated hazard ratios were adjusted for various medications use such as statins, antihypertensives, antiplatelets, nitrates, antidepressants, nonsteroidal anti-inflammatory drugs, alpha-glucosidate inhibitors, thiazolidinediones., and respiratory drugs, as well as for the presence of renal diseases, respiratory conditions, and cancer.

## DISCUSSION

Our findings highlight SIRT1 as a critical gatekeeper in AS pathophysiology, playing a protective role by mitigating oxidative stress, pro-fibrotic, and pro-calcific processes. Notably, SIRT1 also appears to be a mediator in the multifaceted benefits of SGLT2i. Indeed, our results show that SGLT2i reduces primary VICs calcification *in vitro* through endothelial cell-mediated mechanisms, suggesting a novel therapeutic avenue against valve calcification. These findings are further reinforced by real-world data, which highlight the SGLT2i potential to influence AS progression beyond glycemic control, reducing the hospitalization for non-rheumatic aortic valve disease.

Several cellular and molecular pathways contribute to the pathophysiology of AS^16^. However, while the mechanisms driving valvular calcification have been extensively studied, relatively few investigations have focused on identifying protective factors^17^. Thus, in this context, our results confirm the presence of well-known pathways that are dysregulated in AS tissues. They further show the pivotal role of SIRT1 as a key driver of the pathology when down-regulated, offering new insight into its protective potential during AS progression. Even if it has been shown that SIRT1 is down-regulated in AS, its contribution to pathogenesis of this pathology is still poorly investigated^8^. Our data address this knowledge-gap by showing that in AS tissues, genes known to activate SIRT1 are down-regulated^19^ (*e.g.*, *PRKAA*), while its suppressors^20,21^ (*e.g.*, *CTSB*; *CD28*) show an increased expression. Similarly, transcription factors that either promote (*e.g.*, *MYC*; *FOXO3*) or inhibit^22^ (*e.g.*, *HIC1*) SIRT1 expression are regulated in parallel, further highlighting the dysregulation of SIRT1 in AS pathophysiology. Moreover, it is known that SIRT1 plays a fundamental role in several processes, including oxidative stress, inflammation, fibrosis, and calcification. The dysregulation of these processes leads to the onset and development of CVD, including AS^26,36,37^. Building on this, our study specifically investigated the expression of genes involved in these processes in stenotic valves, where SIRT1 is down-regulated. Notably, our data reveal that genes positively regulated by SIRT1 (antioxidant genes) are significantly down-regulated in the AS valve. In contrast, genes linked to fibrosis and calcification pathways are up-regulated. These results together with existing literature shed lights on the intricate gene expression patterns linked to SIRT1 dysregulation in AS pathology. Furthermore, when combining the aortic valve transcriptomic results with the *in vitro* model of SIRT1 overexpression, we found that more than 400 genes directly modulated by SIRT1 were concordant, supporting a causal relationship between SIRT1 and fibrocalcific processes. Reinforcing this, the identification of *ADAMTS7*, *LMO4* and *TMEM44*, recently associated with AS^14^, as genes directly modulated by SIRT1 in our *in vitro* iVICs model, underscores the relevance of SIRT1’s protective mechanisms. Furthermore, given the recent findings showing that *SORT1* is strongly linked to AS, since it is up-regulated in stenotic tissues and play a role in calcification processes^38^, the evidence of its down-regulation by SIRT1 suggest that SIRT1 itself may act as a master regulator of aortic valve fibrosis and calcification. Functional assays further validate the proposed anti-fibrotic and anti- calcific role of SIRT1, revealing for the first time that SIRT1 overexpression, in valvular cells, results in the down-regulation of pro-fibrotic proteins (*e.g.* COL1A1 and αSMA). Additionally, this over-expression also inhibits genes and proteins associated with pro-calcific pathways in AS, such as PCSK9^10^, suggesting a new mechanism to counteract valvular calcification, which include PCSK9 inhibition. Consistently with our observation that SIRT1 inhibits pro-calcific pathways, recent studies reinforce the therapeutic potential of SIRT1 in modulating PCSK9 and its associated pathways. Specifically, it was shown that pharmacological activation of SIRT1 reduces PCSK9 secretion from murine hepatocytes *in vitro* and lowered plasma levels of PCSK9 in atherosclerosis-prone *Apoe^-^/^-^ mice in vivo*^39^. Furthermore, recent preliminary findings evidenced a direct interaction between SIRT1 and PCSK9, wherein SIRT1 inhibits PCSK9 activity through deacetylation, providing a potential mechanism for the interplay between these two proteins^40^.

These molecular findings prompted us to explore whether SIRT1 might also influence AS through a specific genetic predisposition. Although no significant variants within the *SIRT1 locus* were identified as directly associated with AS in the largest GWAS available^14^, the observed association of common SIRT1 variants with cardiovascular traits, particularly SBP, offers critical insight. Indeed, elevated SBP is considered one of the major risk factors for AS^41^, suggesting that SIRT1 may contribute to AS indirectly by modulating blood pressure regulation. This underscores the multifaceted role of SIRT1 in cardiovascular pathology and the importance of considering genetic and molecular mechanisms when unraveling its contribution to AS pathogenesis.

Recently, SGLT2i, have gained prominence for their remarkable cardiovascular benefits, among which their role in blood pressure control stands out^28,42^. Furthermore, of particular interest for our study, SGLT2i enhance the activation of the SIRT1 signaling pathway, positioning them as plausible therapeutic candidates in the AS context^12^. Within the aortic valve, the endothelium plays a pivotal role, covering the valve leaflets, regulateing molecular exchange, and absorption from the bloodstream but also communicating with the underlying layers that house VICs, the main driver of valvular fibro-calcification^43^. Considering this intricate interplay and aiming to address the complexity of the valve, our findings contribute to the understanding of VECs-VICs cell cross-talk. Specifically, we demonstrated that treating the valvular endothelium with SGLT2i not only reshapes their cellular landscape by enhancing the protective subpopulation of VECs, but also modulates VICs calcification. This modulation occurs, at least in part through the secretion of key factors such as DKK1 which in turn inhibits calcification by suppressing Wnt signaling, and ADM, which subsequently promotes NO production, a critical molecule for vascular health^44,45^. Notably, this effect is tightly linked to the activation of the NO-SIRT1 axis by SGLT2i, a pathway essential for endothelial health and calcification inhibition^46^. These findings are consistent with previously proposed mechanisms but are observed here, for the first time, within the specific context of AS in human cells, thereby integrating seamlessly into a broader understanding of valve biology and disease pathophysiology.

Despite extensive research, drug therapies aimed at reducing AS have largely been ineffective. Trials involving statins and other medications have failed to demonstrate significant benefits in slowing the progression of aortic valve calcification or improving clinical outcomes, especially when administered in AS late stages^47^. However, promising avenues are emerging, such as therapies targeting PCSK9 and lipoprotein(a) [Lp(a)], known causal risk factors for both atherosclerosis and AS^10,48^. Emerging treatments, including neutralizing antibodies, small interfering RNAs, and antisense oligonucleotides, have shown potential in lowering PCSK9 and Lp(a) levels^47^. Also, renin-angiotensin-aldosterone system inhibitors have shown potential benefits in reducing pro-fibrotic processes^49^ and the ARBAS (Angiotensin Receptor Blockers in Aortic Stenosis) randomized control trial (https://clinicaltrials.gov/study/NCT04913870) will unveil their effects in AS context. Furthermore, nitric oxide-cyclic guanosine monophosphate signaling pathways are being investigated, with preclinical studies showing that phosphodiesterase type 5 inhibitors and soluble guanylate cyclase activators like ataciguat may reduce aortic valve calcification and slow disease progression (<u>NCT02481258</u>)^47^. Our current real-world analysis highlights the potential protective effect of SGLT2i against aortic valve degeneration. Indeed, our data indicate a significant reduction in the incidence of hospitalization rate for non-rheumatic aortic valve disease among patients treated with SGLT2i compared to those receiving sulfonylureas. This protective effect, consistent across various subgroups and adjusted for comorbidities and concomitant therapies, suggests a promising clinical impact of SGLT2i on AS. However, results from robust randomized controlled trials are still needed to confirm these findings.

In conclusion, our findings underline the therapeutic potential of SGLT2i in AS, highlighting their ability to modulate SIRT1-related pathways and endothelial interactions. By restoring the protective role of SIRT1, this class of cardio-metabolic drugs may have an important impact on slowing the AS progression, reducing the incidence of hospitalization for non-rheumatic aortic valve disease and offering a new strategy for the clinical management of patients affected by this debilitating disease.

